# P bodies coat germ granules to promote transgenerational gene silencing in *C. elegans*

**DOI:** 10.1101/2022.11.01.514641

**Authors:** Zhenzhen Du, Kun Shi, Jordan S. Brown, Tao He, Wei-Sheng Wu, Ying Zhang, Heng-Chi Lee, Donglei Zhang

**Author notes:** These authors contribute equally (Zhenzhen Du, Kun Shi). Co-corresponding author: Heng-Chi Lee and Donglei Zhang.

## Abstract

The formation of biomolecular condensates has emerged as a critical mechanism for compartmentation in living cells. Despite interactions between distinct condensates having been reported, the biological relevance of these interactions remains elusive. In germ cells, small RNA silencing factors are enriched in germ granule condensates, where distinct factors are organized into sub-compartments with specific functions linked to genome surveillance or transgenerational gene silencing. Here we showed that perinuclear germ granules are coated by P body condensates, which are known for housing translationally-inactive mRNAs and mRNA degradation factors. Disruption of P body factors, including CGH-1/DDX6 and CAR-1/LSM14, lead to dispersal of small RNA factors from perinuclear germ granules and disorganization of sub-compartments within germ granules. We further found that CAR-1 promotes the interaction between CGH-1 and germ granule factors, and these interactions are critical for the ability of CGH-1 to promote piRNA-mediated gene silencing. Importantly, we observed that *cgh-1* mutants are competent in triggering gene silencing but exhibit defects in maintaining gene silencing in subsequent generations. Small RNA sequencing further showed that *cgh-1* mutants exhibit defects in amplifying secondary small RNAs, known carriers of gene silencing memories. Together, our results uncover the function of P body factors in small RNA-mediated transgenerational gene silencing and highlight how the formation and function of one condensate can be regulated by an adjacent, interacting condensate in cells.

## Introduction

Formation of biomolecular condensates has emerged as a critical mechanism for membraneless compartmentation in living cells (Banani et al., 2017). The formation of various condensates has been recently linked to diverse functions, including chromatin regulation, RNAi inheritance, RNA processing, cellular signaling. It has also been reported that distinct condensates interact in cells, including Cajal bodies with their attached B-snurposome, Cajal bodies with nucleoli, and Processing bodies (P bodies) with stress granules (Gall et al., 1999; Pena et al., 2001; Sanders et al., 2020). However, little is known about whether these interactions play a role in regulating the formation or function of their cognate interacting condensates.

It has been reported that germ granules associate with P bodies in the early embryos of *C. elegans* and other animals (Gallo et al., 2008; Kotaja et al., 2006; Lin et al., 2006). Germ granules and P bodies are two distinct condensates associated with different biological functions. In germ cells of diverse animals, many small RNA pathway factors are enriched in germ granules (Lim and Kai, 2007; Ouyang and Seydoux, 2022). In *C. elegans*, distinct sub-compartments of germ granules have been reported to house factors linked to distinct functions in the RNAi pathway. Specifically, P granules are enriched for Argonaute family proteins that are critical for RNA surveillance (Batista et al., 2008; Chen et al., 2022; Claycomb et al., 2009a; Gu et al., 2009). Z granules are enriched for factors involved in the transmission of RNAi silencing over generations, termed RNAi inheritance (Ishidate et al., 2018a; Wan et al., 2018). Mutator granules are enriched for factors that are involved in silencing small RNA amplification (Phillips et al., 2012). It has been proposed that factors in these distinct sub-compartments of germ granules coordinate to control gene silencing and its inheritance (Dodson and Kennedy, 2020). P bodies in contrast are well-known as condensates housing mRNA degradation factors and translationally-inactive mRNAs (Sheth and Parker, 2003). Studies have shown that P bodies in the germline and early embryo carry out distinct functions (Boag et al., 2008; Noble et al., 2008), and some P bodies form perinuclear foci in the adult germline where perinuclear germ granules are found (Boag et al., 2008). These observations prompted us to investigate whether P bodies physically interacts with perinuclear germ granules, and if so, to investigate the functional relevance of those interactions.

Here we show that in the *C. elegans* germline, P bodies are situated specifically at the cytoplasmic side of perinuclear germ granules. We found that P body factor CGH-1 and CAR-1, the homologue of human DDX6 and LSM14, interact with germ granule components, particularly with P granule and Z granule factors. Our analyses showed that CAR-1 promotes the formation of CGH-1 condensates and CGH-1’s interaction with germ granule factors. Notably, both CAR-1 and CGH-1 regulate the formation and organization of germ granules, and promote piRNA-mediated gene silencing. Specifically, we showed that CGH-1 plays a critical role in transgenerational gene silencing by promoting the amplification of secondary small RNAs.

## Results

### P bodies are localized to the cytoplasmic side of perinuclear P granules

As described above, previous studies have shown that some P body condensates are enriched at the nuclear periphery in the adult germline of *C. elegans* (Boag et al., 2008). Indeed, by monitoring the localization of different P body factors including CGH-1, LET-711 and IFET-1 using endogenously-tagged fluorescent marker expressing strains, we found that P body condensates present as perinuclear foci in nearly all stages of the adult germline (Figure S1A and S1B). When we simultaneously monitored two P body markers, we found that they are largely co-localized (Figure S1B), showing P body factors co-localize at nuclear periphery. To examine the relationship between P bodies and perinuclear P granules in the adult germline, we simultaneously monitored the localization of P body and P granule markers. We found that P body markers, including CGH-1, LET-711 and IFET-1, are localized to the cytoplasmic side of perinuclear P granule markers, including PGL-1, PRG-1 and CSR-1 (Figure 1 and Figure S1C). When we also monitored the localization of NPP-9 (a nuclear pore marker), we observed that P bodies are situated to the cytoplasmic side of P granules, which together are situated on top of nuclear pore clusters (Figure 1B). To further characterize the relationship between P bodies and adjacent germ granules, we identified and defined individual P bodies as the center and measured the vertical (outside/inside) and horizontal (left/right) distance between individual P body markers to their adjacent germ granule markers (Figure 1C). Using these analyses, we confirmed that the majority of CGH-1 condensates are located directly to the cytoplasmic side of PGL-1 or CSR-1 condensates (Figure 1C). In addition, we noticed that some of the P body and P granule markers exhibit overlapping signals (Figure 1A), indicating these two cellular bodies are partially co-localized. We then examined the relationship between P bodies and Z granules by simultaneously monitoring the localization of CGH-1 and ZNFX-1 or WAGO-4 (two Z granule markers). We found that while most CGH-1 condensates are located slightly toward the cytoplasmic side of Z granules, and they are frequently found as adjacent condensates as indicated by higher variance in horizontal distance between these two condensates (Figure 1C, D). We also observed some partial co-localization of CGH-1 condensates and WAGO-4 condensates (Figure 1D). We then monitored the relationship between P bodies and Mutator foci, and we found that CGH-1 condensates also tend to localize slightly toward the cytoplasmic side and are adjacent to Mutator marker MUT-16 (Figure 1C). Very little co-localizing signal between CGH-1 and MUT-16 is found (Figure S1D). Together with the previous report that PZM granules form tri-condensate assemblages (Wan et al., 2018), our image analyses suggest a spatial organization of P bodies, P granules, Z granules, and Mutator foci at the nuclear periphery (Figure 1E).

**Figure 1.**
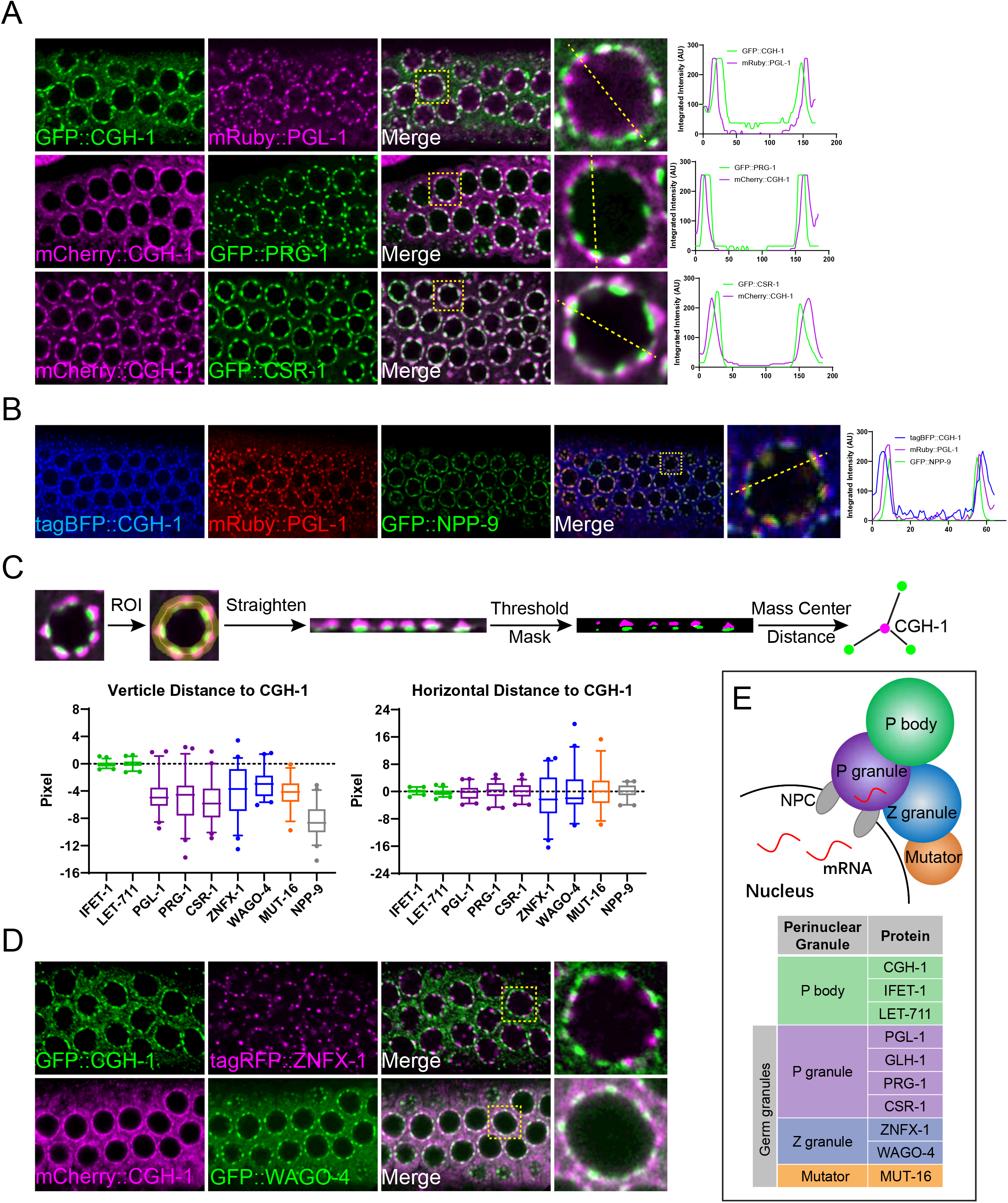
P bodies localize to the cytoplasmic side of germ granules. (a) Fluorescent micrographs show the localization of P body marker CGH-1 and the indicated P granule markers in the pachytene region of adult germlines. The line in the merged image indicates the position of the line scan for measuring fluoresce intensity across single germline nuclei (right). Note that white color foci in the merge image represent the co-localization of two makers. (b) Fluorescent micrographs show the spatial arrangement of P body, P granule and nuclear pore clusters at germline nuclei. The line in the merged image indicates the position of the line scan for measuring fluoresce intensity across single germline nuclei (right). (c) A schematic (top) showing the measurements of vertical distance (bottom left) and horizontal (bottom right) distance between the indicated proteins to P body marker CGH-1 (d) Fluorescent micrographs show the localization of P body marker CGH-1 and the indicated Z granule proteins in the pachytene region of adult germlines. Note that white color foci in the merge image represent the co-localization of two makers. (e) A model depicting spatial arrangements of the P body and distinct sub-compartments of the germ granule, including P granule, Z granule and Mutator foci, at the nuclear periphery.

### CGH-1 interacts with small RNA pathway factors enriched in P granules

Our observations of partial co-localization between perinuclear P body markers and both P granule and Z granule markers raise the possibility that factors between these condensates might interact with each other. To test this hypothesis, we performed mass spectrometry (MS) analyses of CGH-1 complexes (Figure 2A). With CGH-1 Immunoprecipitation (IP) MS analyses, we identified several known P body factors but did not detect factors enriched in P granules or Z granules. To detect more transient interactions, we applied chemical crosslinking before performing IP MS analyses, an approach that has successfully identified transient interactions between ribonucleoprotein complexes (Chen et al., 2022; Ge et al., 2019). Indeed, additional proteins were detected in the crosslinked CGH-1 complexes, including GLH-1 and Argonaute proteins PRG-1, CSR-1 and WAGO-1 (Figure 2A and Figure S2A). The presence of these Argonaute proteins in crosslinked CGH-1 complexes was confirmed by IP Western analyses (Figure 2B). While we did not detect the presence of Z granule factors, such as ZNFX-1 or WAGO-4, in crosslinked CGH-1 complexes, we performed reciprocal IP-MS analyses of crosslinked ZFNX-1 and WAGO-4 complexes and were able to detect the presence of P body factors, specifically CGH-1 and CAR-1 (Figure 2C and 2D). Together, our MS analyses showed that P body factors CAR-1 and CGH-1 interact with the small RNA pathway factors enriched in P granule and Z granule factors. Our observations also raise the possibility that P bodies may regulate the localization or function of P granules and/or Z granules.

**Figure 2.**
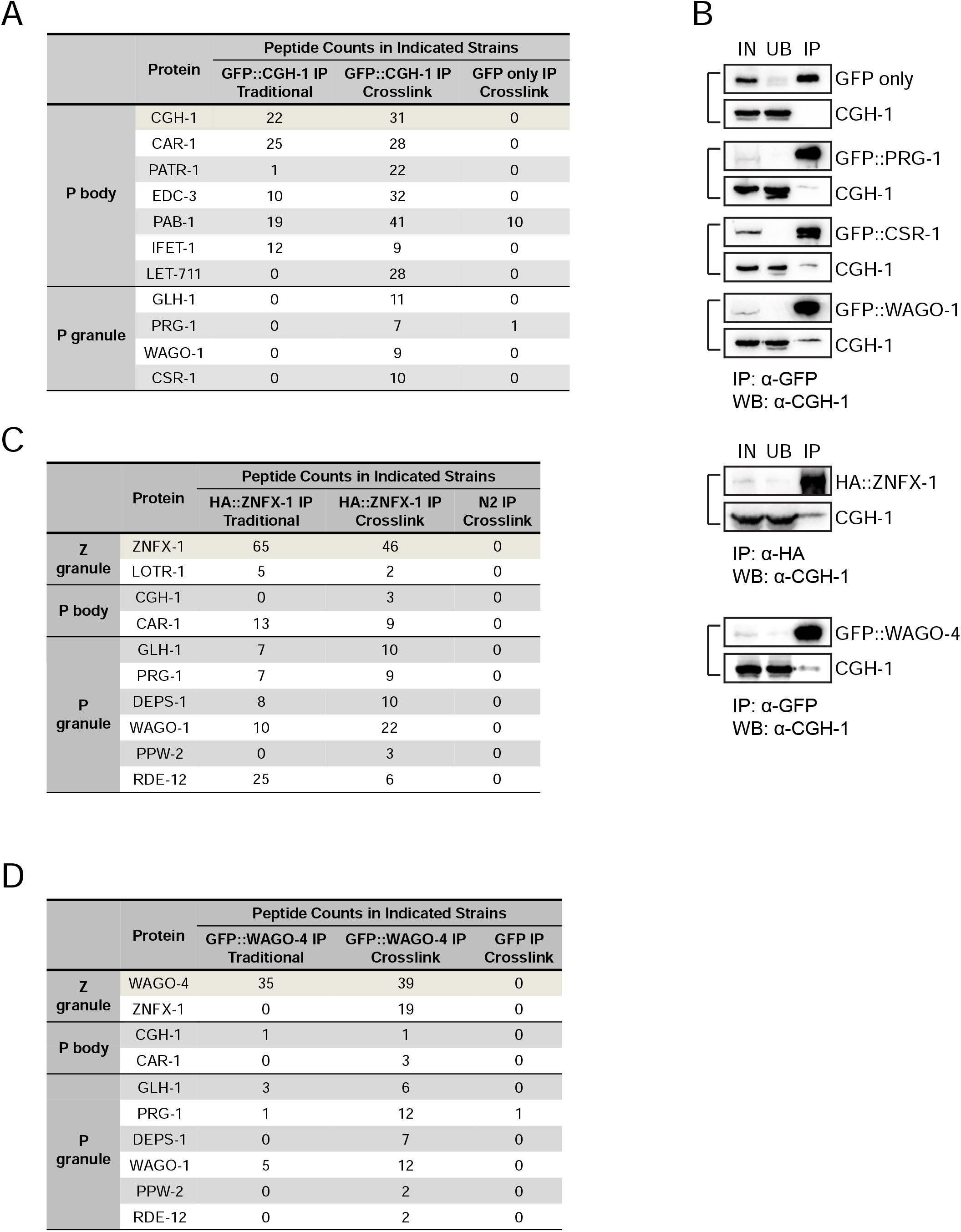
Proteomic analyses of interactions between P body and germ granule factors. (a) Proteomic analyses of CGH-1 complex. The numbers of peptides identified in the crosslinked or non-crosslinked condition are shown. GFP only IP experiment in the crosslink condition served as a control. (b) Western blot analyses show the interactions between CGH-1 and P granule factors PRG-1, WAGO-1 or CSR-1. IN: input, UB: unbound fraction, IP: immunoprecipitated fraction. (c) Proteomic analyses of ZNFX-1 complex. The numbers of peptides identified in the crosslinked or non-crosslinked condition are shown. N2 control IP experiment in the crosslink condition served as a control. (d) Proteomic analyses of WAGO-4 complex. The numbers of peptides identified in the crosslinked or non-crosslinked condition are shown. N2 control IP experiment in the crosslink condition served as a control.

### CGH-1 promotes the localization of small RNA components in germ granules

We then examined whether CGH-1 regulates the localization of small RNA factors in germ granules. We first noticed that *cgh-1* partial deletion mutant (*ok492*) animals exhibit a germline nuclei organization defect, where some germline nuclei detach from the surface and move to the center of the gonad (Figure 3A). Importantly, we found that the perinuclear localization of both PRG-1 and CSR-1 were compromised in *cgh-1* mutants, including in the partial deletion mutant (*ok492*) or a temperature-sensitive mutant (*tn691*) (Figure 3A top, Figure S3A, S3B). Fluorescent recovery after photobleaching (FRAP) analyses further indicated that CGH-1 promotes the turnover of PRG-1 and CSR-1 in P granules (Figure S3C). We then examined the role of CGH-1 in regulating Z granules and Mutator foci. We found that the perinuclear localization of Z granule factors, including ZNFX-1 and WAGO-4, as well as the perinuclear localization of MUT-16, are also compromised in *cgh-1* mutants (Figure 3B, 3C, and S3D-F). Interestingly, the localization of P granule assembly factors, including PGL-1 and GLH-1, are not affected in *cgh-1 (ok492)* or *cgh-1 (tn691)* mutants (Figure 3A bottom, Figure S3A, S3B). Together, our imaging analyses suggest that CGH-1 specifically promotes the localization of various small RNA pathway factors to germ granules.

**Figure 3.**
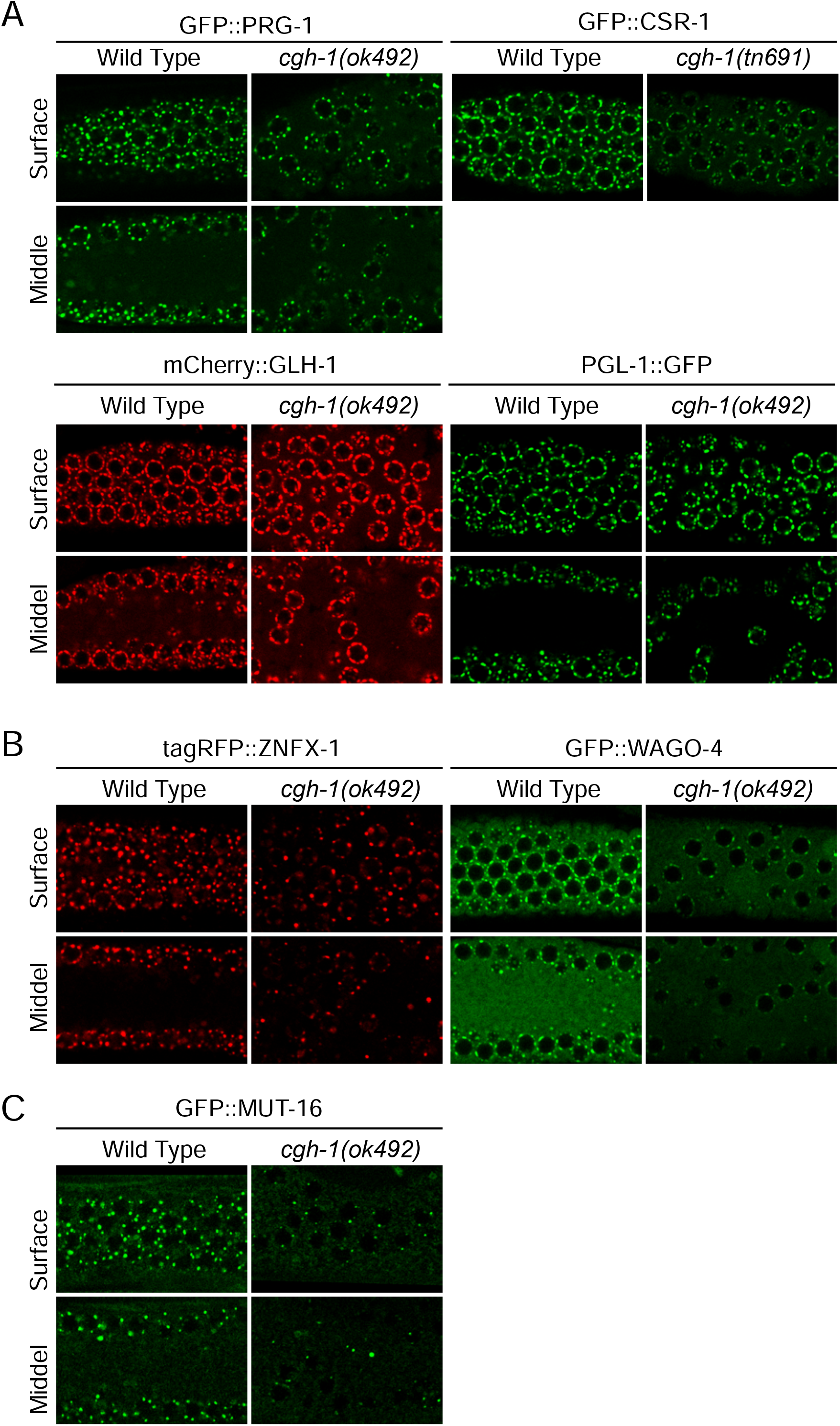
P body factor CGH-1 promotes the localization of small RNA pathway factors at perinuclear germ granules. Fluorescent micrographs show the localization of indicated P granule markers (A), Z granule markers (B) and Mutator foci marker MUT-16 (C) in the pachytene region of adult germlines in wild type or in *cgh-1 (ok492)* mutant animals. The images from surface or middle section of the germline are shown.

We then examined whether the localization of P body factor CGH-1 to perinuclear foci requires P granule assembly. VASA-like helicases GLH-1 and GLH-4 have been reported to promote the perinuclear localization of various P granule factors (Chen et al., 2020; Spike et al., 2008). Surprisingly, while the localization of P granule factors, such as PRG-1, become diffuse upon *glh-1* and *glh-4* double RNAi treatment, the localization of CGH-1 is not affected (Figure S3G). In addition, we observed that CGH-1 perinuclear localization is not affected in mutants of the PRG-1 pathway, including *prg-1*, *hrde-1*, *znfx-1* or *mut-16* mutants (Figure S3H). This indicates that CGH-1’s perinuclear localization does not rely on P granule assembly factors GLH-1 and GLH-4 or piRNA pathway factors and implies that an alternative pathway contributes to the perinuclear localization of CGH-1.

### CAR-1 promotes the interaction of CGH-1 with germ granules and supports CGH-1’s role in piRNA silencing

Our recent study has suggested that P granule localization of small RNA pathway factors is critical for the proper recognition of mRNA targets by piRNAs (Chen et al., 2022). As the perinuclear localization of small RNA pathway factors is also compromised in *cgh-1* mutants, we wondered whether piRNA-mediated gene silencing is defective in *cgh-1* mutants. We used two sensitized piRNA reporters where the silencing of these reporters depends on both factors required for initiation and maintenance of piRNA silencing (Chen et al., 2022; Cordeiro Rodrigues et al., 2019). We found that silencing of these piRNA reporters is compromised in *cgh-1 (tn691)* mutant animals, *cgh-1* (*dz407*) mutant animals, and in *cgh-1* RNAi-treated animals (Figure 4A, 4B and Figure S4A, S4B). We then examined the roles of other P body factors in piRNA silencing. We found that only RNAi of *car-1*, but not other P body factors examined, led to the activation of our piRNA reporter (Figure 4C). Remarkably, only RNAi knockdown of *car-1*, but not that other P body factors examined, led to the dispersal of CGH-1 from perinuclear P bodies to the cytoplasm (Figure 4D). The levels of CGH-1 in *car-1* RNAi treatments were not changed (Figure S4C). These observations show that CAR-1 promotes the formation of CGH-1 condensates and raises the possibility that CGH-1’s perinuclear localization may be critical for CGH-1’s interaction with small RNA factors and for its function in piRNA silencing. We therefore examined whether the interactions between CGH-1 with P granule factors require CAR-1. Indeed, the interactions between CGH-1 with P granule factors WAGO-1 and PRG-1 were greatly reduced in *car-1* RNAi treated animals (Figure 4E). Similar to *cgh-1* mutants, the perinuclear localization of PRG-1 and WAGO-4 are also reduced in *car-1* RNAi treated animals (Figure S4D). These observations show that the ability for CGH-1 to form perinuclear condensates correlates with CGH-1’s ability to interact with germ granule factors and to regulate germ granule formation. Together, our reporter analyses suggest that CGH-1 and its perinuclear localization are critical for piRNA-mediated gene silencing.

**Figure 4.**
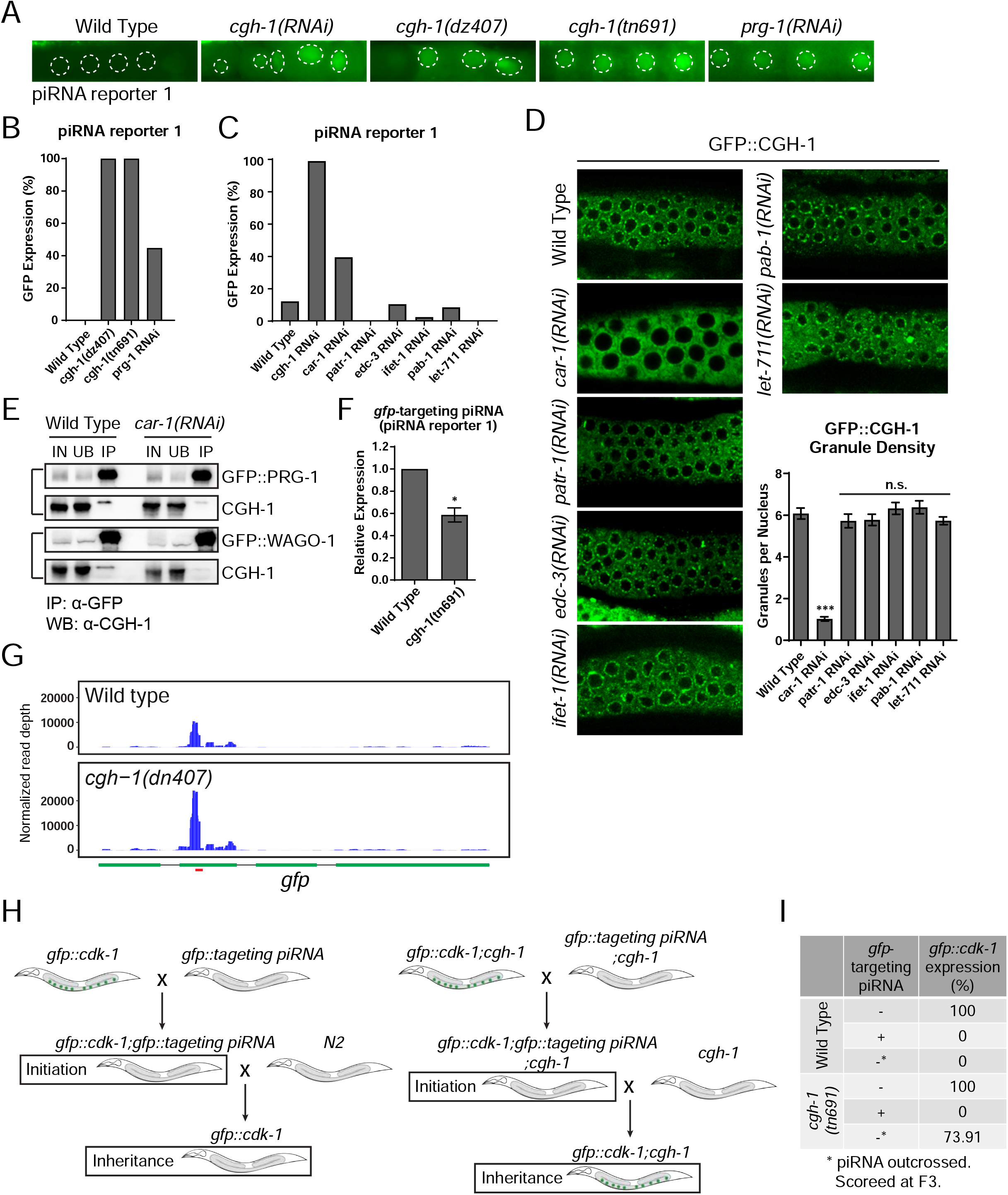
P body factor CGH-1 and CAR-1 promote piRNA-dependent gene silencing. (a) GFP expression of the piRNA reporter #1 in the indicated strains. Dotted circles indicate the location of maturing oocyte nuclei. (b) Percentage of screened animals showing GFP expression in the piRNA reporter #1 in the indicated strains. (c) Percentage of screened animals showing GFP expression in the piRNA reporter #1 in the indicated strains. (d) Fluorescent micrographs show the localization of P body marker CGH-1 the indicated RNAi-treated animals. Granule density (number of granule/nuclei/section) of GFP∷CGH-1 in wild type or the indicated RNAi-treated animals was calculated. These quantifications correspond to micrographs in Figure 4D). Statistical analysis was performed using a one-tailed Student’s t-test. Bars indicate the mean, errors bars indicate the standard deviation. (e) Western blot analyses showing that the interaction between CGH-1 and P granule factors PRG-1 or WAGO-1 is reduced in *car-1* RNAi treated animals. IN: input, UB: unbound fraction, IP: immune-precipitated fraction (f) q-RT PCR measurements of the abundance of the GFP-targeting piRNA in the indicated strains. Statistical analysis was performed using a two-tailed Student’s t-test. Bars indicate the mean, errors bars indicate the standard deviation for 3 technical replicates from each genotype. (g) 22G-RNAs distribution at GFP coding sequences in the indicated strains. The red bar (below) indicates the location of the GFP sequence complementary to the GFP-targeting piRNA. (h) A schematic showing the GFP reporter assays to determine the role of CGH-1 in the initiation or maintenance of piRNA silencing. (i) Percentage of GFP reporter expression from screened animals in the indicated strains. Note that *cgh-1(tn691)* mutant animals can initiate piRNA-directed silencing of the GFP reporter but exhibit defects in the inheritance of gene silencing.

### CGH-1 promotes the inheritance of piRNA silencing

piRNAs trigger gene silencing against their targets through the production of secondary 22G-RNAs that are loaded onto WAGO Argonautes to mediate gene silencing (Batista et al., 2008; Lee et al., 2012). To examine how CGH-1 contributes to piRNA silencing, we compared the piRNA and downstream 22G-RNA levels in wild type versus *cgh-1* mutants. We observed a reduced accumulation of anti-GFP synthetic piRNA but surprisingly increased levels of secondary 22 RNAs in *cgh-1* mutants at the synthetic piRNA targeting site (Figure 4F and 4G). These results suggest that CGH-1 is not required for the biogenesis of the synthetic piRNA or for the production of 22G RNAs around the piRNA targeting site. It has been previously shown that piRNAs can trigger transgenerational gene silencing in *C. elegans* (Ashe et al., 2012; Lee et al., 2012; Shirayama et al., 2012). As described above, we found that CGH-1 is present in ZNFX-1 and WAGO-4 complexes, both factors required for transgenerational RNAi (Ishidate et al., 2018b; Wan et al., 2018; Xu et al., 2018), and CGH-1 promotes ZNFX-1 and WAGO-4 localization in Z granules. In addition, we showed that the sensitized piRNA reporter is activated following RNAi knockdown of *znfx-1* (Figure S4B). Together, these observations imply that CGH-1 could play a regulatory role in piRNA silencing, perhaps by promoting the inheritance of piRNA-induced gene silencing.

As CGH-1 is an essential gene, we cannot examine the role of CGH-1 in transgenerational gene silencing using a *cgh-1* null allele. We reasoned that if the function of *cgh-1* in the temperature-sensitive mutant (*tn691*) is partially compromised when these animals are grown at the permissive temperature, then this mutant would allow us to examine the role of CGH-1 in transgenerational gene silencing. In support of this hypothesis, we noticed that *cgh-1* (*tn691*) mutant animals grown at the permissive temperature (20 degree *Celsius*) already exhibit a more mild but similar phenotype to those grown at the non-permissive temperature (25 degree *Celsius*), such as significantly reduced broods and the compromised formation of ZNFX-1 granules (Figure S5). We then examined the role of CGH-1 in the initiation of piRNA-induced silencing by crossing a strain carrying a synthetic GFP-targeting piRNA to the strain carrying a GFP reporter (Figure 4H). We found that the synthetic GFP-targeting piRNA can trigger the silencing of the GFP reporter in both wild type and in *cgh-1* (*tn691*) mutant animals (Figure 4I) grown at 20 degree *Celsius*. This result suggests that *cgh-1* (*tn691*) mutant animals are competent for triggering piRNA-directed gene silencing when raised at the permissive temperature. To examine the role of CGH-1 in the maintenance of gene silencing memory, we outcrossed the synthetic GFP-targeting piRNA from the wild type or *cgh-1* (*tn691*) mutant background and found that while all wild type animals continue to exhibit silencing of the GFP reporter in the F3 generation, over 73% of *cgh-1* (*tn691*) mutant animals exhibit expression (re-activation) of the GFP reporter (Figure 4I). These observations suggest that CGH-1 plays a critical role to promote transgenerational gene silencing triggered by piRNAs.

### CGH-1 promotes RNAi inheritance

Similar to piRNAs, double stranded RNA (dsRNA) can also trigger transgenerational gene silencing lasting for several generations, a phenomenon also known as RNAi inheritance (Alcazar et al., 2008). To determine whether CGH-1 plays a role in RNAi inheritance, we treated wild type and *cgh-1* (*tn691*) mutants with GFP RNAi food (bacteria expressing dsRNA targeting GFP) and monitored the expression of the GFP reporter over generations (Figure 5A). We found that in the RNAi treated generation (P0), both wild type and *cgh-1 (tn691)* mutant animals exhibited robust GFP silencing, suggesting both strains are competent in triggering dsRNA-induced gene silencing (Figure 5B and Figure 5C). Remarkably, while over 70% of the wild type animals continue to keep the GFP reporter silenced for over three generations, less than 30% of the *cgh-1* mutants can maintain GFP silencing for a single generation in the absence of dsRNA (Figure 5B and 5C). These observations suggest that CGH-1 promotes transgenerational gene silencing triggered by both piRNA and dsRNA.

**Figure 5.**
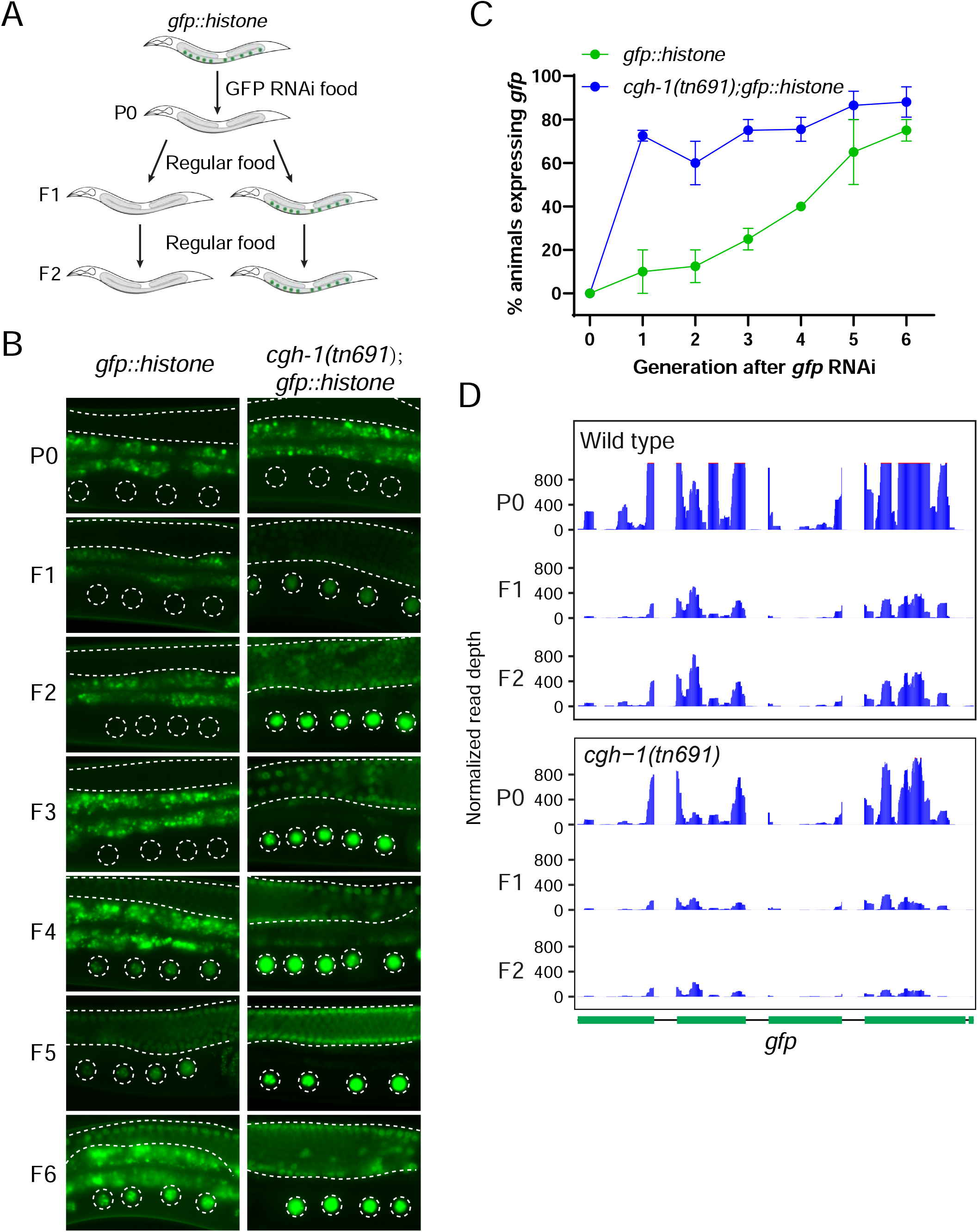
CGH-1 promotes RNAi inheritance and 22G-RNA synthesis in inheriting generations. (a) A schematic showing the RNAi assay with a GFP∷histone reporter to determine whether the strain can initiate RNAi and inherit gene silencing over generations. In these experiments, both wild type and *cgh-1 (tn691)* mutant animals are grown at the permissive temperature (20 degree *Celsius*). (b) Fluorescent micrographs show representative images of the GFP∷H2B reporter in the RNAi-treated generation (P0) or the following generations (F1 to F6) in wild type and *cgh-1 (tn691)* mutant animals (c) Percentage of screened animals showing GFP reporter expression in wild type and *cgh-1 (tn691)* mutant animals (d) Antisense 22G-RNAs distribution mapped to GFP coding sequences in the indicated strains from the RNAi-treated animals (P0) or the following generations (F1 or F2).

As previous studies have demonstrated the role of 22G-RNAs in RNAi inheritance (Ouyang et al., 2022; Xu et al., 2018), we performed small RNA sequencing to measure the levels of 22G-RNAs in the wild type and in the *cgh-1* (*tn691*) mutants in the parental RNAi treated animals as well as the untreated inheriting F1 and F2 progeny. In wild type animals, we found that the production of anti-sense GFP targeting 22G-RNAs are highest in the RNAi-treated generation (P0) then accumulation is reduced in the F1 and F2 generations in comparison to RNAi treated animals (Figure 5D). Notably, levels of anti-sense GFP targeting 22G-RNAs were reduced in the RNAi-treated animals (P0) of *cgh-1* (*tn691*) mutant animals and were further reduced in F1 and F2 generations compared to wild type animals. Together, our data indicate that CGH-1 promotes the amplification of 22G-RNAs and thus is critical for RNAi inheritance.

### CGH-1 promotes the organization of germ granules into distinct sub-compartments

As described above, *C. elegans* germ granules are organized into distinct subcompartments, including P granules, Z granules and mutator foci (Wan et al., 2018). As P granules, Z granules and Mutator foci are enriched for factors involved in genome surveillance, RNAi inheritance and 22G-RNA production, we wondered whether CGH-1 may regulate the organization of PZM granules to promote RNAi-inheritance and 22G-RNA synthesis. We first confirmed that P granule marker PGL-1 and Z granule marker ZNFX-1 are partially separated in in the adult germline, consistent with the previous report that these two granules are organized to distinct but partially overlapped subcompartment (Figure 6A) (Wan et al., 2018). Remarkably, we found that the separation of the Z and P granule markers is compromised in *cgh-1 (tn691)* mutants and in *cgh-1* RNAi treated animals, where they are nearly all co-localized (Figure 6A and 6B). The co-localization between PGL-1 and ZNFX-1 is also increased in *car-1* RNAi treated animals compared to control RNAi treated animals (Figure 6A and 6B). These observations suggest that CGH-1 and CAR-1 not only promote the formation of phase separated condensates of RNAi factors (Figure 3A-C), but also promote the proper organization of germ granules.

**Figure 6.**
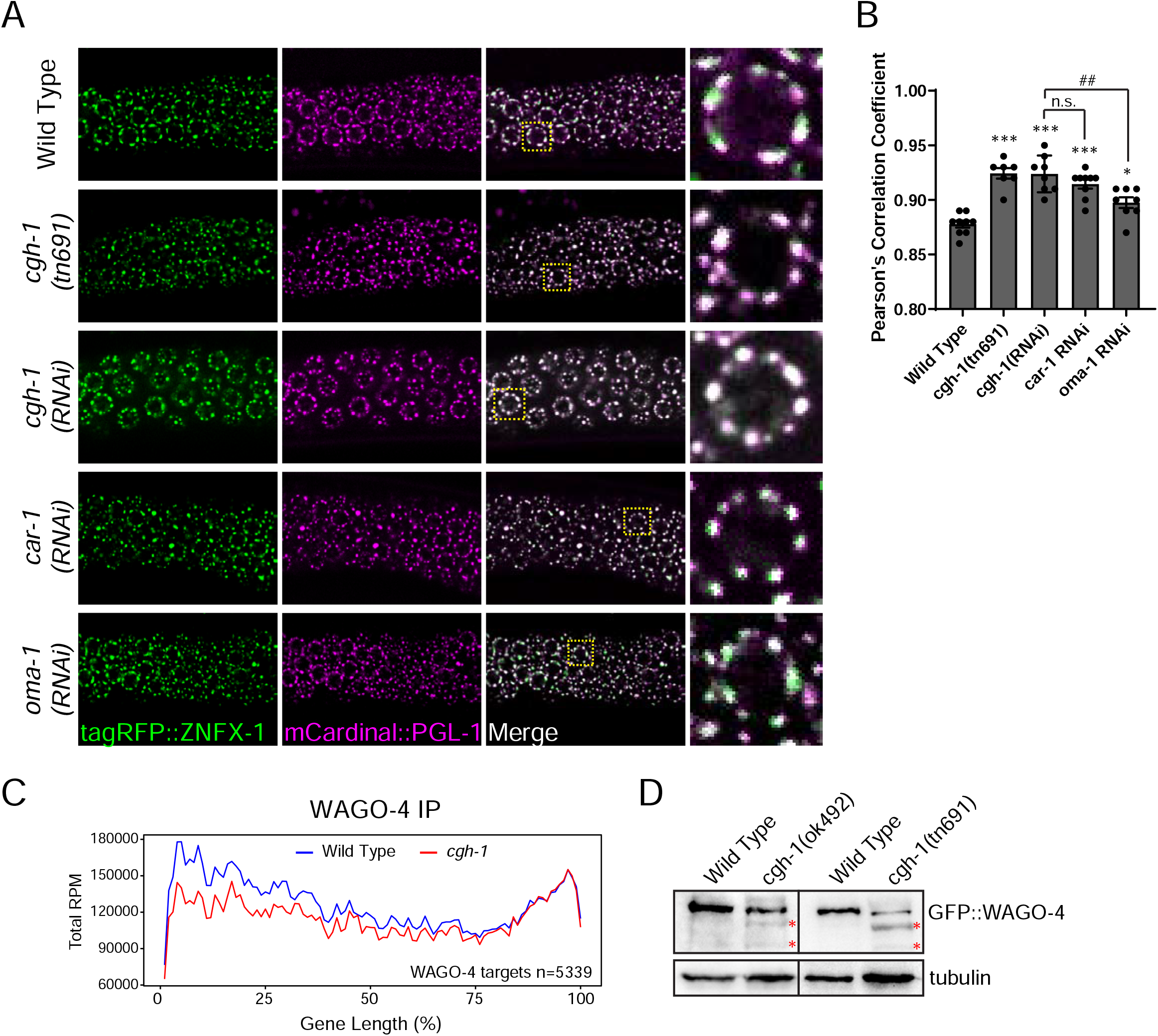
CGH-1 promotes proper germ granule organization and WAGO-4 22G-RNA production. (a) Fluorescent micrographs show the localization of P granule marker PGL-1 and Z granule marker ZNFX-1 in the indicated mutant or RNAi-treated animals. (b) Pearson’s correlation coefficient of PGL-1 and ZNFX-1 signals in the indicated mutant or RNAi-treated animals. (c) Metagene traces show the total accumulation of WAGO-4 associated 22G-RNAs by percentage of WAGO-4 target gene length in IP experiments. Reads from wild type IP experiments are traced in blue and reads from *cgh-1* RNAi-treated IP experiments are traced in red. (d) Western blots show the levels of WAGO-4 in wild type or in the indicated *cgh-1 mutants*. The red asterisks indicated

### CGH-1 promotes the stability of WAGO-4 Argonaute and the level of its associated small RNAs

Previous studies have shown that WAGO-4 localize to Z granules and play a critical role in RNAi inheritance (Wan et al., 2018; Xu et al., 2018). As we found that CGH-1 is required for RNAi inheritance and promotes the organization of P and Z granules, one interesting hypothesis is that CGH-1 and CAR-1 regulate the trafficking of mRNA/protein complexes between P and Z granules so that 22G-RNAs can be made and associated with WAGO-4 to trigger transgenerational gene silencing. If so, we would expect the accumulation of WAGO-4 22G-RNAs to be compromised in CGH-1 mutants. To test this hypothesis, we performed WAGO-4 small RNA sequencing in wild type and *cgh-1* RNAi treated animals. Since previous studies have shown that several mutants exhibit a shift in 22G-RNA accumulation on mRNAs, we also monitored the distribution of 22G-RNAs. We noticed that the production of WAGO-4 22G RNAs is reduced in *cgh-1* RNAi treated animals, especially at the 5’ ends of mRNAs (Figure 6C). On the contrary, the production of CSR-1 22G-RNAs is slightly increased in *cgh-1* RNAi treated animals, especially at the 3’ end of mRNAs (Figure 6SA). Furthermore, while the levels of PRG-1, CRS-1 and GLH-1 protein remained normal, WAGO-4 levels were reduced in *cgh-1* mutants and WAGO-4 appears to be partially degraded (Figure 6D and S6B). As the absence of small RNAs can trigger the degradation of their associated Argonaute (Choudhary et al., 2007), the aberrant degradation of WAGO-4 proteins in *cgh-1* mutants (Figure 3C) may be caused by the reduction in WAGO-4 small RNAs levels. Together, our observations shows the critical roles of CGH-1 in producing WAGO-4 small RNAs and further support a model where CGH-1 promotes transgenerational gene silencing by facilitating the production of WAGO-4 22G-RNAs (Figure 7).

**Figure 7.**
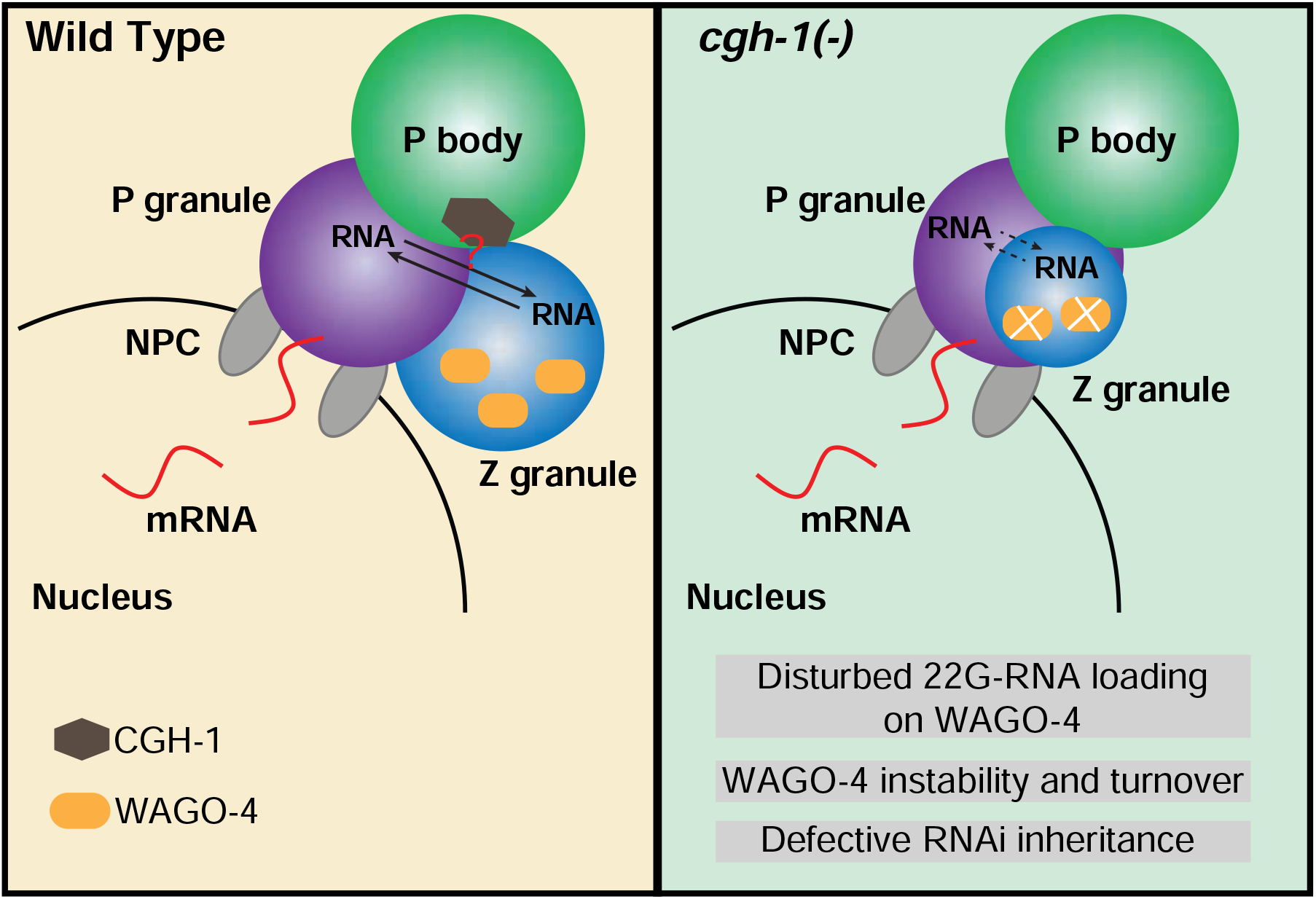
A model showing the roles of P body factor CGH-1 in germ granule organization and WAGO-4 22G-RNA production that promote transgenerational gene silencing.

## Discussion

P bodies and germ granules are well-known cellular condensates, playing important roles in storage of translationally inactive mRNAs and small RNA-based gene silencing, respectively. In this study, we showed that these two distinct phase-separated condensates interact in the germline in *C. elegans* and this interaction is critical for germ granule formation and for transmitting gene silencing signals across generations.

Specifically, our imaging analyses revealed the spatial relationship between P bodies and perinuclear germ granules, where P bodies are situated cytoplasmic to germ granules and nuclear pore clusters (Figure 1B). Using Mass Spectrometry and IP-western analyses, we confirmed the physical interactions between P body factors (specifically CGH-1 and CAR-1) and P/Z granule factors. Our analyses further revealed that P body factors, including CGH-1 and CAR-1, promote the formation and proper organization of germ granules. Interestingly, CGH-1 and CAR-1 only promote the phase separation of small RNA pathway factors, such as PRG-1, CSR-1 and WAGO-4, but not of germ granule assembly factors, such as PGL-1 or GLH-1 (Figure 3A and Figure S4D). In addition, in *cgh-1* and *car-1* depleted animals the organization of distinct P and Z granules within germ granules is compromised, where these granules are aberrantly co-localized (Figure 6A). These are intriguing findings as small RNA factors enriched in P granules and Z granules are linked to functions in mRNA surveillance and RNAi inheritance, respectively. In addition, our analyses suggest that while CGH-1 regulates the formation and organization of germ granules, germ granule assembly is not essential for the formation of perinuclear CGH-1 condensates (Figure S3H). This result suggests that CGH-1 can be recruited to perinuclear foci through a GLH-1/GLH-4-independent mechanism.

Importantly, we further describe the functional significance of P body - germ granule interaction. We showed that piRNA silencing is compromised in both *cgh-1* mutants and *car-1* depleted animals. We further observed that CAR-1 is critical for the perinuclear localization of CGH-1 (Figure 4D), and that loss of CAR-1 compromises the interaction between CGH-1 and P granule factors (Figure 4E). Together, our data support a model where the interactions between P bodies and germ granules are critical for CGH-1 to regulate small RNA-mediated gene silencing.

Our genetic analyses further showed that CGH-1 is not required for the initiation of gene silencing triggered by either piRNAs or dsRNA, but rather is critical for the transmission of gene silencing across generations. Defects in secondary small RNA accumulation further suggest a critical role for CGH-1 in promoting small RNA amplification to promote transgenerational gene silencing.

Our transcriptome-wide analyses of secondary small RNAs associated with WAGO-4 suggest that CGH-1 is required for the efficient production of WAGO-4 22G-RNAs. Interestingly, it has been proposed that CGH-1 homologue DDX6 may regulate stress granule formation by promoting mRNA transfer from P bodies to stress granules (Hondele et al., 2019). It is currently unclear whether the mis-organization of P/Z granules observed in *cgh-1* mutants is the cause or consequence of the defect in small RNA amplification. Nonetheless, these observations suggest that CGH-1 and CAR-1 may specifically contribute to the trafficking of mRNPs involved in gene silencing. One intriguing model is that CGH-1 is critical for the transference of silenced mRNA from P granules to Z granules which will act as a template for small RNA amplification and enable WAGO-4 22G-RNA loading, critical for transgenerational gene silencing.

In summary, our study uncovers the critical roles of P body factors in the formation of germ granules and their function in transgenerational gene silencing. It will be interesting to examine whether germ granules play a reciprocal role in regulating P body function. As more cases of interactions between distinct cellular condensates are reported, our study highlights an exciting example where the formation, organization, and function of one condensate can be controlled by factors from an adjacent, interacting condensate.

## Methods

### *C. elegans* strains

*C. elegans* were maintained on standard nematode growth media (NGM) plates seeded with the Escherichia coli OP50 strain at 20°C or temperatures where indicated. Strains and alleles used in this study are listed in Supplementary Table 1. Some strains were obtained from the Caenorhabditis Genetics Center (CGC).

### RNA interference

RNA interference (RNAi) was performed by feeding animals with E. coli HT115 (DE3) strains expressing the appropriate double-stranded RNA (dsRNA). RNAi bacterial strains were obtained from the Ahringer *C. elegans* RNAi Collection (Source BioScience) or the collection derived from *C. elegans* ORFeome Library (Horizon Discovery). Bacterial cμltures were grown in Luria broth supplemented with 100 μg/ml ampicillin overnight at 37°C. Cultures were seeded on NGM plates containing 100 μg/ml ampicillin and 1 mM IPTG and incubated at room temperature for 24 hours. L1 or L4 hermaphrodites were picked onto the plates for feeding at 20°C prior to score for phenotypes. HT115 (DE3) expressing empty RNAi vector L4440 was used as the control.

### Brood Size

Single hermaphrodite L4s (P0s) were placed onto individual freshly seeded NGM plates and allowed to grow for 24 hours at 16, 20 or 25°. P0 adults were transferred to new NGM plates every 24 hours until they no longer laid eggs. All the F1 progeny on each plate were counted. The brood size of each P0 animal is the total sum of F1s for all plates where the P0 animal laid eggs. At least 10 P0 animals were used to calculate the brood size for each strain.

### CRISPR

CRISPR-Cas9 guide RNAs were designed with the online IDT Alt-R Custom Cas9 crRNA Design Tool. DNA fragments were amplified by PCR and transcribed to sgRNAs in vitro with HiScribe Quick T7 High Yield RNA Synthesis Kit (New England Biolabs), and sgRNAs were purified with Monarch RNA Cleanup Kit (New England Biolabs). sgRNA sequences and PCR primers used in this study are listed in Supplementary Table 2.

Partially single-stranded DNAs was used as donor templates for in situ gene knock-in using a modified method from (Dokshin et al., 2018). GFP/mCherry/tagBFP coding sequences and the coding sequences with 40 bp homology arms were amplified by PCR. Purified PCR products were mixed as 1:1 and treated by a denaturation and renaturation process [95°C 2:00 minutes, 85°C 10 seconds, 75°C 10 seconds, 65°C 10 seconds, 55°C 1:00 minute, 45° 30 seconds, 35° 10 seconds, 25° 10 seconds, 4° hold] to generate hybrid DNA donors.

CRISPR experiments were conducted with Cas9 RNP strategies. Final concentrations of injection components were as follows: 250 ng/μL of Alt-R Cas9 protein (IDT) and 200 ng/uL in vitro synthesized sgRNA. All components were mixed and incubated at 37°C for 10 min, then 200 ng/μL of hybrid DNA donor and 40 ng/μL of pRF4 *rol-6 (dm)* were added. Injection mixtures were introduced into the germline of *C. elegans* young adults by microinjection. The F1 rollers were isolated and genotyped to identify the desired transgenes.

### Imaging

Fluorescent proteins were visualized in living nematodes by mounting young adult animals on 2% agarose pads with M9 buffer [22 mM KH_2_PO_4_, 42 mM Na_2_HPO_4_, and 86 mM NaCl] with 10-50 mM levamisole. Fluorescent images were captured using a Zeiss LSM800 confocal microscope equipped with AiryScan detector and a Plan-Apochromat 63X/1.4 Oil DIC M27 objective. Colocalization studies between CGH-1 and P granule proteins were performed by the 2D Superresolution mode for Zeiss Airyscan. Images were taken by ZEN 2.3 (blue edition) acquisition software.

Images were processed and germ granule fluorescence was quantified in ImageJ. For germ granule density analysis, 10 nuclei in each germline pachytene region were randomly selected and perinuclear puncta were counted manually. For quantitative co-localization analysis, all image manipulations were performed with ImageJ using the Coloc 2 plugin. Images of 8-12 gonads were collected and quantified.

### Fluorescence recovery after photobleaching

Photobleaching and fluorescence recovery of GFP∷PRG-1 and GFP∷CSR-1 were conducted and recorded using a Zeiss LSM 800 confocal microscope with a Plan-Apochromat 63×/1.4 Oil DIC M27 objective, controlled by the ZEN software. The ZEN built-in fluorescence recovery after photobleaching (FRAP) module was used to perform FRAP experiments. Bleaching was performed using 100% laser power in the 488nm channel. Multiple granules were selected in each imaging field and fluorescence intensity of selected granules was bleached to less than 20% of the intensity before photobleaching. Images were acquired every 1 second during a recovery phase of 120 seconds after bleaching. Images were processed and quantified in ImageJ. The total fluorescence intensity was measured for areas containing bleached granules (I), unbleached granules (Inorm), or areas without granules (Ibkg) at each time point. Fluorescence recovery ratios were calculated as (In−Ibkgn)(Inormi/Inormn)/(Ii−Ibkgi), where n stands for time points after photobleaching and i stands for initial phase before photobleaching. Recovery ratios for >10 granules at different time points were calculated for each strain.

### Immunoprecipitation

A total of ~100,000 synchronized young adult animals were used for immunoprecipitation. Synchronized animals were washed with M9 buffer three times before frozen in liquid nitrogen and stored at −80°C. Pellets were resuspended in equal volumes of immunoprecipitation buffer [20 mM Tris-HCl pH 7.5, 150 mM NaCl, 2.5 mM MgCl_2_, 0.5% NP-40, 80 U ml^−1^ RNase inhibitor (Invitrogen), 1 mM dithiothreitol, and protease inhibitor cocktail without EDTA (Promega)], and grinded in a glass grinder for 8-10 times. The whole grinding process should not exceed 10 minutes. Lysates were clarified by spinning down at 15000 rpm, 4°C, for 15 minutes. Supernatants were incubated with GFP-Trap magnetic agarose beads (ChromoTek) or anti-HA-tag magnetic beads (MBL) at 4°C for 2 hours. Beads were washed with IP wash buffer [20 mM Tris–HCl pH 7.5, 150 mM NaCl, 2.5 mM MgCl_2_, 0.5% NP-40, and 1 mM dithiothreitol] six times for 10 minutes each time and PBS buffer twice for 5 minutes each time, and then resuspended in PBS buffer for mass spectrometry analysis, western blot, or RNA extraction.

### Chemical crosslink

Chemical cross-linking of proteins was performed with dithio-bismaleimidoethane (DTME) (Sigma-Aldrich) dissolved in dimethyl sulfoxide (DMSO). ~100,000 synchronized young adults were collected and washed three times with M9 buffer. M9 buffer was discarded to the same amount of the worm volume, then DTME was added to a final concentration of 2 mM. Samples were incubated for 30 minutes at room temperature with occasional shaking then washed three times with M9 buffer to remove excess cross-linker. Worm pellets were resuspended in equal volume of immunoprecipitation buffer [20 mM Tri-HCl pH 7.5, 150 mM NaCl, 2.5 mM MgCl_2_, 0.5% NP-40, 0.5 mM dithiothreitol, 80 U/mL RNase inhibitor (Invitrogen), protease inhibitor cocktail without EDTA (Promega)]. Worm pellets were homogenized using glass homogenizer for 15-20 strokes on ice. Lysates were centrifuged at 14000 x g for 10 minutes to remove insoluble material. Supernatants were incubated with GFP-Trap magnetic agarose beads (ChromoTek) or anti-HA-tag magnetic beads (MBL) for 2 hours at 4°C on an end-to-end rotator. Supernatant was removed and beads were washed with 1 mL of wash buffer [20 mM Tris-HCl pH 7.5, 150 mM NaCl, 2.5 mM MgCl_2_, 0.5% NP-40] six times for 10 minutes each time and with a final wash of 0.05% NP-40. Beads were incubated at 37°C for 30 minutes in 50 μL de-crosslinking buffer [50 mM Tris-HCl pH 7.5, 150 mM NaCl, 2 mM MgCl_2_, 0.2% Tween 20, 10 mM dithiothreitol]. The final samples were boiled in 2 x SDS loading buffer at 100°C for 5 minutes, then analyzed by mass spectrometry.

### Mass spectrometry

The samples were subjected to SDS PAGE gel electrophoresis experiments, and gel slices were subjected to in-gel digestion with Trypsin (Promega) at 37°C overnight. Purified peptides were desalted using a C18 Stage Tip column (Pierce), concentrated and dried. Then the peptides are reconstituted with 0.1% formic acid aqueous solution for subsequent liquid chromatography with tandem mass spectrometry (LC-MS/MS) analysis using a Q Exactive HF-X mass spectrometer (Thermo Scientific) equipped with a nanoliter flow rate Easy-nLC 1200 chromatography system (Thermo Scientific). The buffers used in the liquid chromatography separation consisted of buffer A [0.1% formic acid] and buffer B [80% acetonitrile, 0.1% formic acid]. The liquid chromatography separation was achieved by linear gradient elution at a flow rate of 300 nL/min. The related liquid phase gradient was as follows: 0-3 minutes, linear gradient of liquid B from 2% to 8%; 3-43 minutes, linear gradient of liquid B from 8% to 28%; 43-51 minutes, linear gradient of liquid B from 28% to 40%; 51-52 minutes, linear gradient of liquid B from 40% to 100%; 52-60 minutes, linear gradient of liquid B maintained at 100%. The Q Exactive HF-X mass spectrometer was used in the data dependent acquisition (DDA) mode. The full scan survey mass spectrometry analysis was performed as detection mode of positive ion, analysis time of 60 minutes, scan range of 350-1800 m/z, resolution of 60,000 at m/z 20, AGC target of 3e6, and maximum injection time (IT) of 50 ms. The 20 highest intensity precursor ions were analyzed by the secondary mass spectrometry (MS2) with resolution of 15,000 at m/z 200, AGC target of 1e5, and maximum IT of 25 ms, activation type of HCD, isolation window of 1.6 m/z, and normalized collision energy of 28. The mass spectrometry data were analyzed using MaxQuant 1.6.1.0 and used the database from Uniprot Protein Database (species: *Caenorhabditis elegans*).

### Western blot

Immunoprecipitated proteins or lysates prepared from ~100 synchronized young adult worms by boiling worms in worm boiling buffer [100 Mm Tris-HCl pH 6.8, 8% SDS, 20 mM 2-mercaptoethanol] for 10 minutes and in 1× SDS sample buffer for 5 minutes were used for western blot. Proteins were separated by standard SDS-PAGE and transferred to PVDF membranes (Roche) using a Trans-Blot Turbo Transfer System (Bio-Rad). Membranes were blocked in TBST (20 mM Tris-HCl pH 7.4, 150 mM NaCl, and 0.05% Tween-20) containing 5% skimmed milk for 2 hours at room temperature, then probed with primary antibodies against CGH-1 (1:2000), GFP (1:500, Santa Cruz Biotechnology), mCherry (1:1000, Abcam), HA (1:1000, Cell Signaling Technology) or tubulin (1:2000, Sigma-Aldrich) overnight at 4°C. After five washes with TBST buffer, the membranes were incubated with secondary antibodies against the species of the primary antibodies at room temperature for 1 hour and then washed five times with TBST. ECL substrates were used for detection of proteins by a Tanon 5200 Chemiluminescent Imaging System (Tanon).

### RNA extraction and small RNA libraries

Total RNA was extracted using the standard method with TRIzol reagent (Invitrogen) from immunoprecipitated samples or whole animals of ~100,000 synchronized young adults.

Small (<200nt) RNAs were enriched with mirVana miRNA Isolation Kit (Ambion). In brief, 80 μL (200-300 μg) of total RNA, 400 μL of mirVana lysis/binding buffer and 48 μL of mirVana homogenate buffer were mixed well and incubated at room temperature for 5 minutes. Then 176 μL of 100% ethanol was added and samples spun at 2500 x g for 4 minutes at room temperature to pellet large (>200nt) RNAs. The supernatant was transferred to a new tube and small (<200nt) RNAs were precipitated with pre-cooled isopropanol at −80°C. Small RNAs were pelleted at 20,000 x g at 4°C for 30 minutes, washed once with 70% pre-cooled ethanol, and dissolved with nuclease-free water. 10 μg of small RNA was fractionated on a 15% PAGE/7M urea gel, and RNA from 18 nt to 30 nt was excised from the gel. RNA was extracted by soaking the gel in 2 gel volumes of NaCl TE buffer [0.3 M NaCl, 10 mM Tris-HCl, 1 mM EDTA, pH 7.5] overnight. The supernatant was collected through a gel filtration column, RNA precipitated with isopropanol, washed once with 70% ethanol, and resuspended with 15 μL nuclease-free water. RNA samples were treated with RppH to convert 22G-RNA 5’ triphosphate to monophosphate in 1x reaction buffer, 10U RppH (New England Biolabs), and 20U RNase inhibitor (Invitrogen) for 3 hours at 37°C, followed by 5 minutes at 65°C to inactivate RppH. RNA was then concentrated with the RNA Clean and Concentrator-5 Kit (Zymo Research). Small RNA libraries were prepared according to the manufacturer’s protocol of the NEBNext Multiplex Small RNA Sample Prep Set for Illumina-Library Preparation (New England Biolabs). The required number of PCR cycles to amplify library is determined by running PCR for 10-15 cycles and analyzing PCR products on a 6% PAGE gel. NEBNext Multiplex Oligos for Illumina Index Primers (New England Biolabs) were used for library preparation. Libraries were sequenced using Illumina NovaSeq 6000 system.

### Bioinformatics

#### sRNA-seq

Fastq reads were trimmed using custom perl scripts. Trimmed reads were aligned to the *C.elegans* genome build WS230 with GFP transgenes added as new chromosomes using bowtie ver 1.2.1.1 (Langmead et al., 2009) with options -v 0 --best --strata. After alignment, reads that were between 17-40 nucleotides in length were overlapped with genomic features (rRNAs, tRNAs, snoRNAs, miRNAs, piRNAs, protein-coding genes, pseudogenes, transposons) using bedtools intersect (Quinlan and Hall, 2010). Sense and antisense reads mapping to individual miRNAs, piRNAs, protein-coding genes, pseudogenes, RNA/DNA transposons, simple repeats, and satellites were totaled and normalized to reads per million (RPM) by multiplying by 1e6 and dividing read counts by total mapped reads, minus reads mapping to structural RNAs (rRNAs, tRNAs, snoRNAs) because these sense reads likely represent degraded products. Reads mapping to multiple loci were penalized by dividing the read count by the number of loci they perfectly aligned to. 22G-RNAs were defined as 21 to 23 nucleotide long reads with a 5’G that aligned to protein-coding genes, pseudogenes, or transposons. RPM values were then used in all downstream analyses using custom R scripts using R version 4.0.3 (R Core Team, 2020), which rely on packages ggplot2 (Wickham, 2016), reshape2 (Wickham, 2007), ggpubr (Kassambara, 2020), dplyr (Wickham et al., 2021). Browser images were constructed by calculating normalized read depth using bedtools genomecov (Quinlan and Hall, 2010) to obtain bedgraph files for each library, then a custom R script was used to build plots.

#### Metagene analysis

Metagene profiles across gene lengths were calculated by computing the depth at each genomic position using 21 to 23 nucleotide long small RNA reads with a 5’G using bedtools genomecov (Quinlan and Hall, 2010). A custom R script was then used to divide genes into 100 bins and sum the normalized depth within each bin. Groups of genes were then plotted using the sum of the normalized depth at each bin. Traces represent normalized depth for 22G-RNAs mapping to CSR-1 targeted genes (n=4932), defined as genes whose mapped 22G-RNAs exhibit over two-fold enrichment from CSR-1 IP than that from input 22G-RNAs (Claycomb et al., 2009b) or WAGO-4 targeted genes (n=5339), defined as genes with at least 5 22G-RNA reads per million mapped reads in IP and input libraries, whose mapped 22G-RNAs exhibit over two-fold enrichment from CSR-1 IP than that from input 22G-RNAs averaged between two biological replicates.

### Stem-loop real-time PCR

Stem-loop real-time PCR was performed to measure piRNA levels. 1 μg of total RNA was reverse transcribed with HiScriptII Q Select RT SuperMix for qPCR (Vazyme) and 50 pM stem-loop RT primer 5’-CTCAACTGGTGTCGTGGAGTCGGCAATTCAGTTGAG-n8-3’ (n8=reverse complement sequences of last 8 nucleotide acids in piRNAs). Each real-time PCR reaction consisted of a 15 uL mixture including 4 μL of cDNA, 1 μM forward piRNA primer 5’-ACACTCCAGCTGGG-n16-3’ (n16= first 16 nucleotide acids in piRNAs), 1 μM universal reverse primer 5’-CTCAAGTGTCGTGGAGTCGGCAA-3’ and ChamQ SYBR qPCR Master Mix (Vazyme). Real-time PCR was performed on a CFX Connect Thermal Cycler (Bio-Rad). Amplification was performed with a two-step reaction at 95°C for 10 minutes and 40 cycles at 95°C for 15 seconds and at 60°C for 30 seconds. The relative fold changes in related piRNAs were normalized to expression levels of mir-35. Each experiment was repeated three times. The PCR primers used for stem-loop real-time PCR are listed in Supplementary Table 2.

### RNAi inheritance assay

Synchronized L1 animals (P0) were RNAi treated by feeding bacteria expressing *gfp* dsRNA or the empty vector L4440. F1 embryos were collected by bleaching P0 animals and grown on regular OP50 bacteria. Embryos from following generations were collected by the same method. The expression of GFP in ~80 animals in each generation was imaged and scored on a Zeiss Axio Scope A1 with a Plan-Neofluar 63x/1.25 Oil M27 objective and a Retiga R6 charge-coupled device camera. Total RNA from P0, F1 and F2 generations were extracted and small RNAs were cloned and sequenced as described above. All RNAi inheritance assays were performed at 20°C.

---

**Figure S1.**
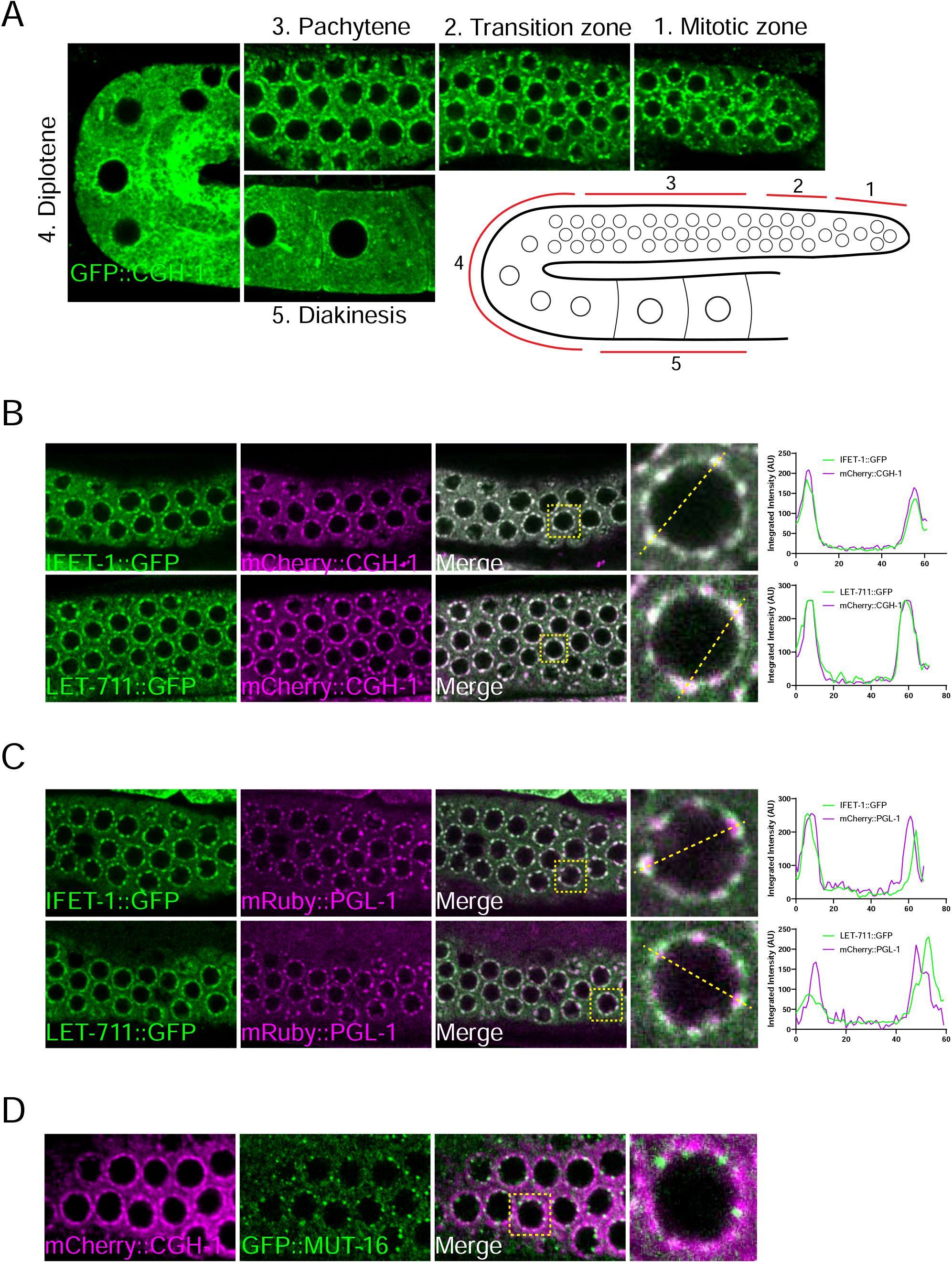
P bodies localize to the cytoplasmic side of germ granules. (a) Fluorescent micrographs show the localization of P body marker CGH-1 in the indicated regions of the *C. elegans* germline. A cartoon shows the relative positions of distinct regions in the gonad of *C. elegans* (bottom right). (b) Fluorescent micrographs show the co-localization of two distinct P body markers in the pachytene region of adult germlines. The line in the merged image indicates the position of the line scan for measuring fluoresce intensity across single germline nuclei (right). (c) Fluorescent micrographs show the localization of the indicated P body and P granule markers in the pachytene region of adult germlines. The line in the merged image indicates the position of the line scan for measuring fluoresce intensity across single germline nuclei (right). (d) Fluorescent micrographs show the localization of the P body marker CGH-1 and Mutator foci marker MUT-16 in the pachytene region of the adult germline.

**Figure S2.**
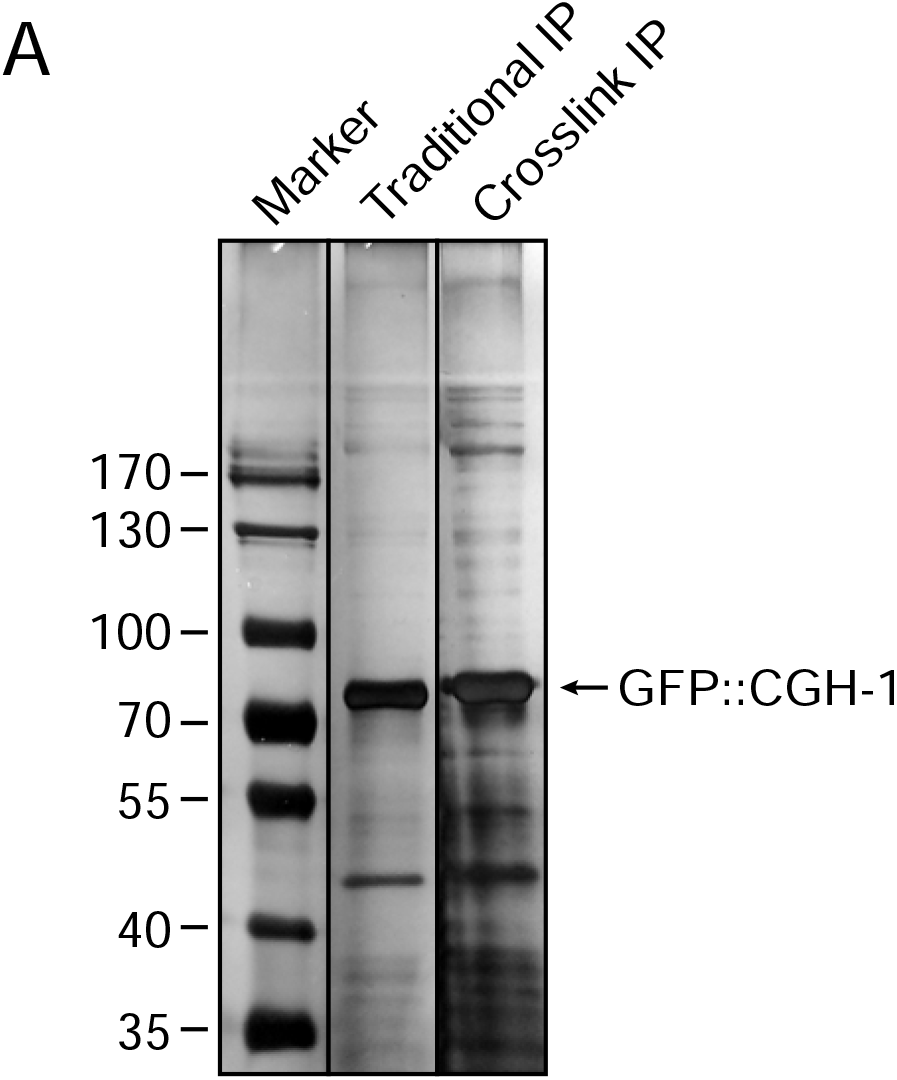
Silver staining of immunoprecipitated CGH-1 complex with or without chemical crosslinking. Note several additional bands were co-immunoprecipitated with CGH-1 when chemical crosslinking was performed before immunoprecipitation.

**Figure S3.**
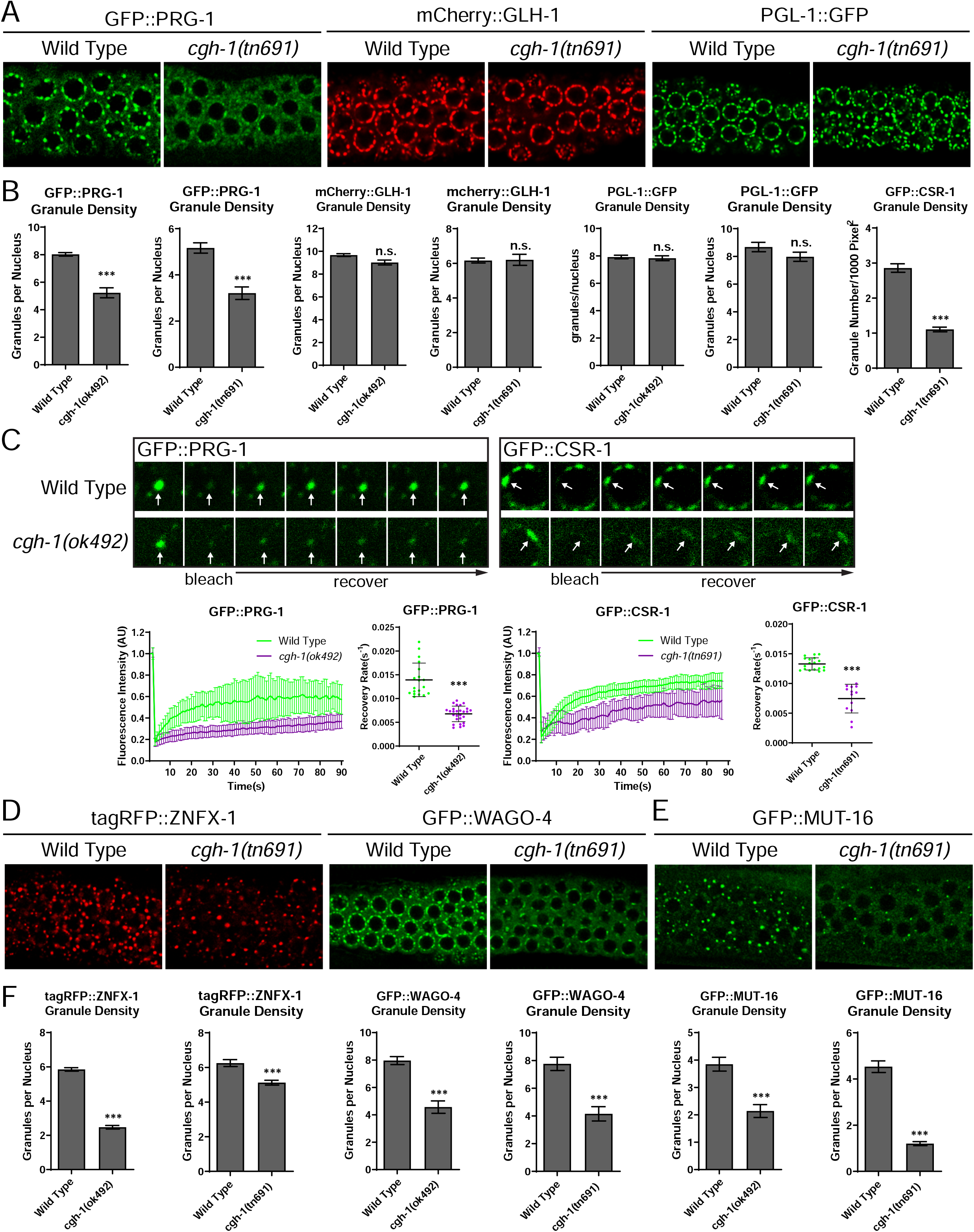
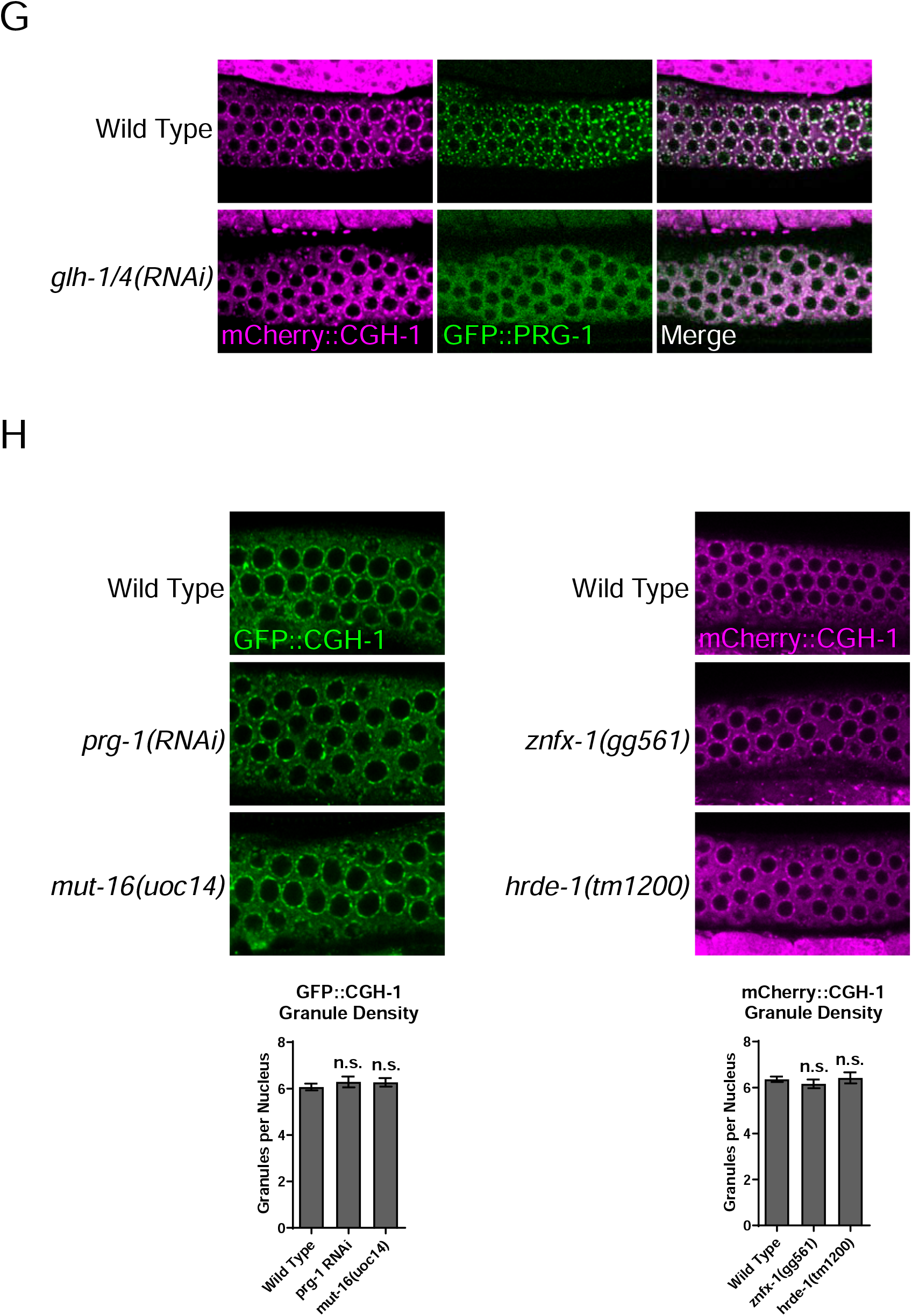
P body factor CGH-1 promotes the localization of small RNA pathway factors at perinuclear germ granules. (a) Fluorescent micrographs show the localization of indicated P granule factors in wild type or in *cgh-1(tn691)* mutant animals. (b) Granule density (number of granule/nuclei/section) of indicated proteins in wild type or the indicated *cgh-1* mutants. These quantifications correspond to micrographs in Figure 3A and Figure S3A). Statistical analysis was performed using a one-tailed Student’s t-test. Bars indicate the mean, errors bars indicate the standard deviation. (c) Fluorescence recovery after photobleaching (FRAP) analyses indicate that the dynamics of indicated P granule factors, including PRG-1 and CSR-1, are reduced in *cgh-1 (ok492)* mutant animals. The arrows indicate the P granules that are photobleached. The quantification of FRAP analyses show the average fluorescence signals of GFP∷PRG-1 or GFP∷CSR-1 at the indicated times (seconds) after photobleaching. Error bars indicate the standard deviations of fluorescence intensities. (d) Fluorescent micrographs show the localization of indicated Z granule markers in the pachytene region of adult germlines in wild type or in *cgh-1 (tn691)* mutant animals. (e) Fluorescent micrographs show the localization of indicated Mutator foci marker MUT-16 in the pachytene region of adult germlines in wild type or *cgh-1 (tn691)* mutant animals. (f) Granule density (number of granule/nuclei/section) of indicated proteins in wild type or the indicated *cgh-1* mutants. These quantifications correspond to micrographs in Figure 3B, 3C and Figure S3D-E). Statistical analysis was performed using a one-tailed Student’s t-test. Bars indicate the mean, errors bars indicate the standard deviation. *(g)* Fluorescent micrographs show the perinuclear localization of P granule factor PRG-1, but not P body factor CGH-1, is reduced in *glh-1* and *glh-4* double RNAi treated animals *(h)* Fluorescent micrographs show the localization of P body marker GFP∷CGH-1 in the pachytene region of adult germlines in wild type or in the indicated mutant or RNAi-treated animals.

**Figure S4.**
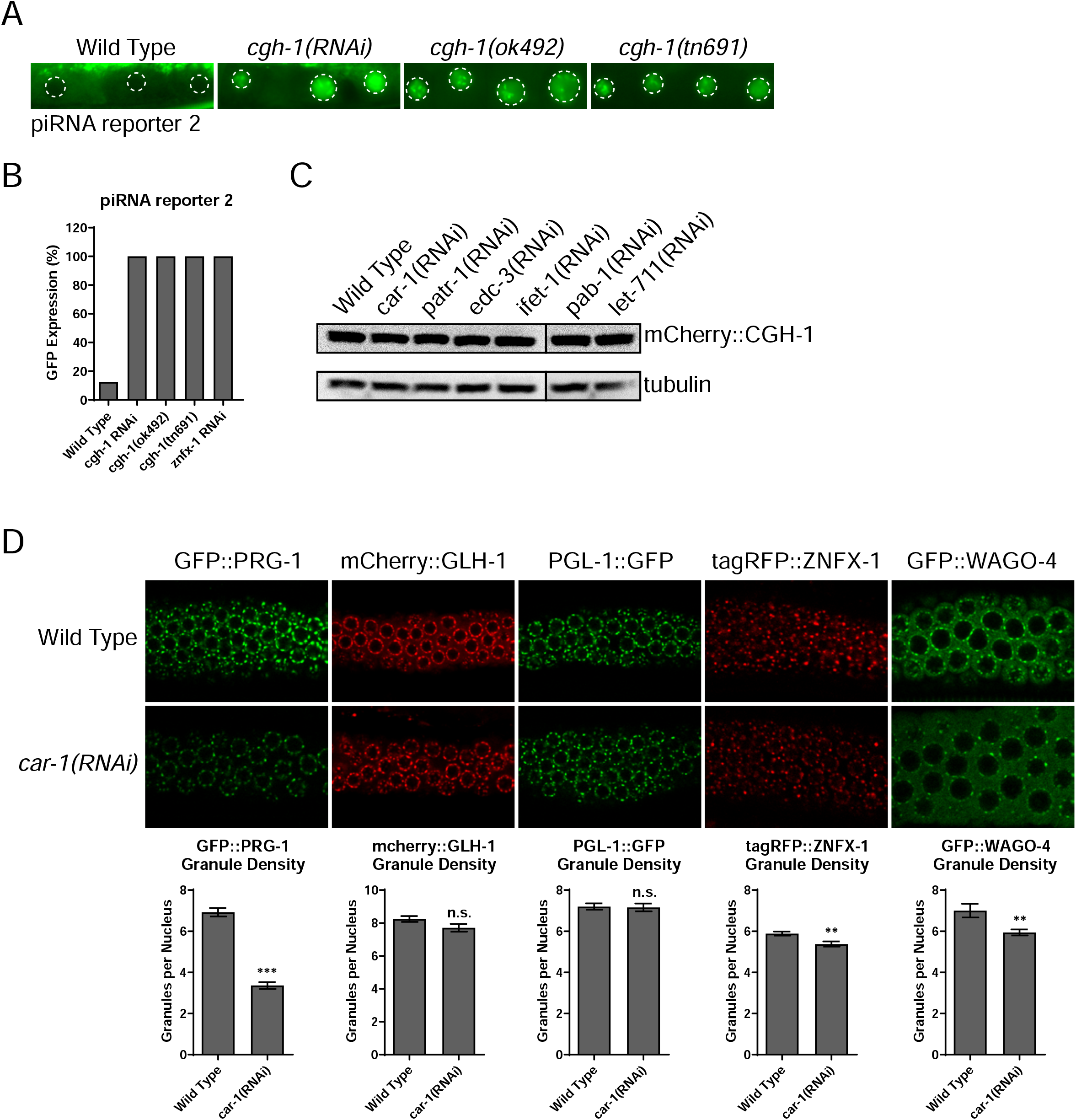
P body factor CGH-1 and CAR-1 are required for piRNA-dependent gene silencing and for the perinuclear localization of small RNA factors. (a) GFP expression in the piRNA reporter #2 in the indicated strains. Dotted circles indicate the location of maturing oocyte nuclei. (b) Percentage of screened animals showing GFP expression in the piRNA reporter #2 in the indicated strains. (c) Western blots show the levels of CGH-1 in the indicated RNAi-treated animals. (d) Fluorescent micrographs show the localization of indicated proteins in the pachytene region of adult germlines in wild type of in *car-1* RNAi-treated animals. Granule density (number of granule/nuclei/section) of indicated proteins in wild type or the *car-1* RNAi-treated animals was calculated. These quantifications correspond to micrographs in Figure S4D. Statistical analysis was performed using a one-tailed Student’s t-test. Bars indicate the mean, errors bars indicate the standard deviation.

**Figure S5.**
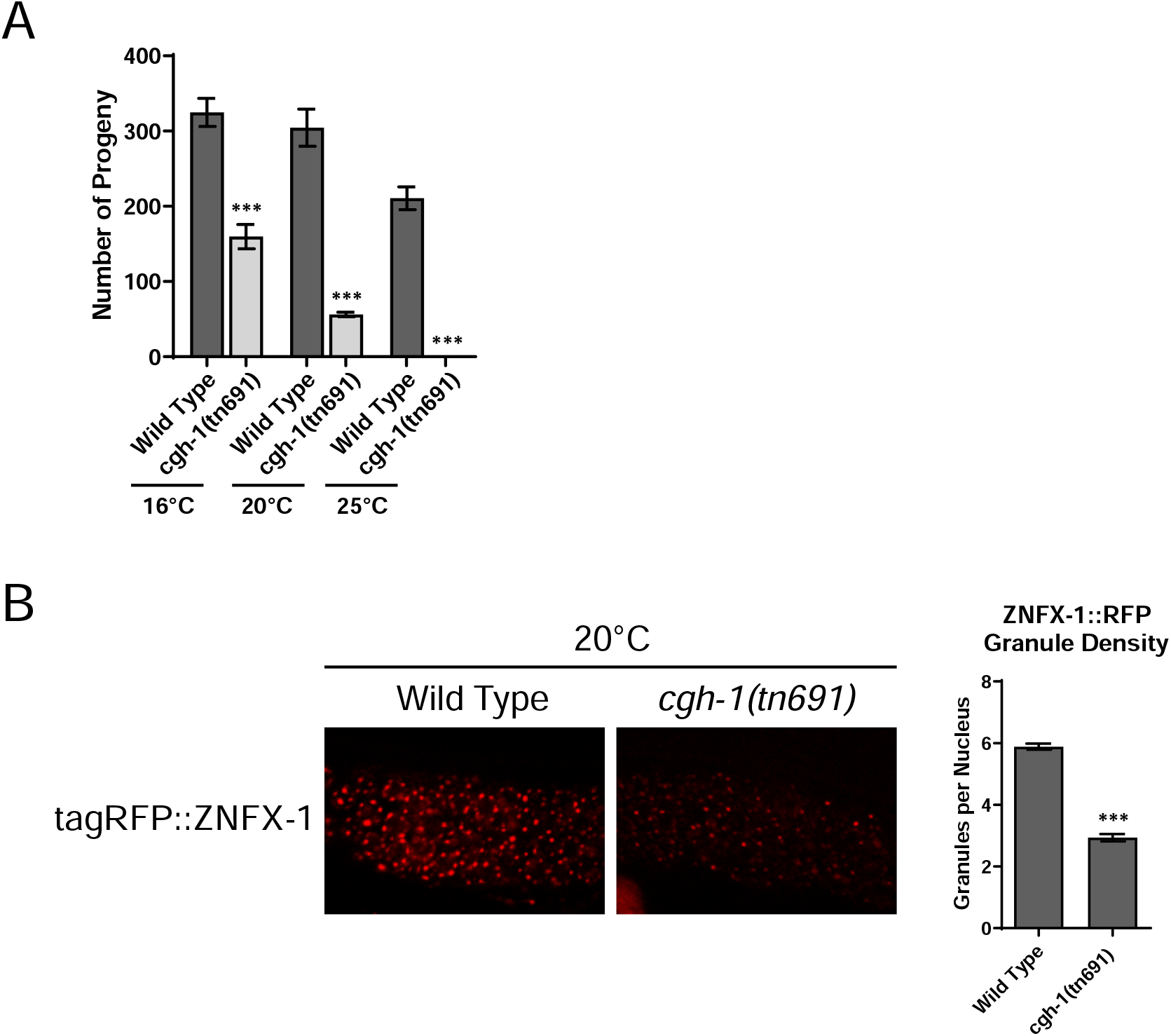
Characterization of temperature-sensitive *cgh-1 (tn691)* strain grown at permissive and non-permissive temperature. (a) The brood size of wild type or *cgh-1 (tn691)* mutant animals grown at the indicated temperature. (b) Fluorescent micrographs show the localization of indicated Z granule factor ZNFX-1 in the pachytene region of adult germlines in wild type or *cgh-1 (ok492)* mutant animals grown at 20 degree Celsius.

**Figure S6.**
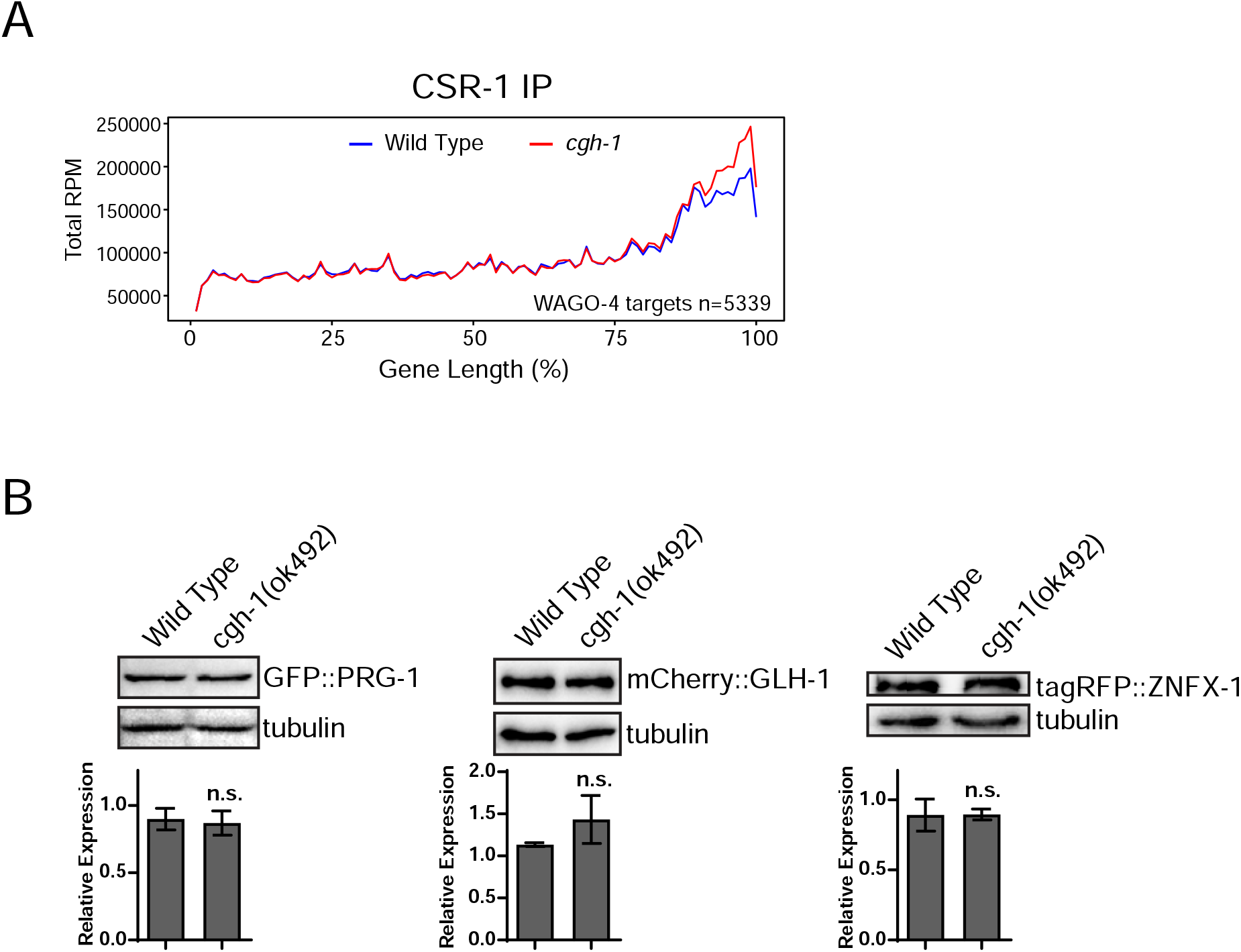
The roles of CGH-1 in small RNA production. (a) Metagene traces show the total accumulation of CSR-1 associated 22G-RNAs by percentage of CSR-1 target gene length in IP experiments. Reads from wild type IP experiments are traced in blue and reads from *cgh-1* RNAi-treated IP experiments are traced in red. (b) Western blots show the levels of indicated proteins in wild type or in the indicated *cgh-1 (ok492) mutants*.

## Notes

### Competing Interest Statement

The authors have declared no competing interest.

